# Run-and-tumble dynamics of *E. coli* is governed by its mechanical properties

**DOI:** 10.1101/2025.01.15.633156

**Authors:** Bohan Wu-Zhang, Peixin Zhang, Renaud Baillou, Anke Lindner, Eric Clément, Gerhard Gompper, Dmitry A. Fedosov

**Affiliations:** Theoretical Physics of Living Matter, Institute for Advanced Simulation, Forschungszentrum Jülich, 52425 Jülich, Germany; PMMH, UMR 7636 CNRS, ESPCI Paris, PSL Research University, Sorbonne Université and Université Paris Cité, 7-9 quai Saint-Bernard, Paris, 75005, France; Institut Universitaire de France, Paris

## Abstract

The huge variety of microorganisms motivates fundamental studies of their behavior with a possibility to construct artificial mimics. A prominent example is the *E. coli* bacterium which employs several helical flagella to exhibit a motility pattern that alternates between run (directional swimming) and tumble (change in swimming direction) phases. We establish a detailed *E. coli* model, coupled to fluid flow described by the dissipative particle dynamics method, and investigate its run-and-tumble behavior. Different *E. coli* characteristics, including body geometry, flagella bending rigidity, the number of flagella and their arrangement at the body are considered. Experiments are also performed to directly compare with the model. Interestingly, in both simulations and experiments, the swimming velocity is nearly independent of the number of flagella. The rigidity of a hook (the short part of a flagellum which connects it directly to the motor), polymorphic transformation (spontaneous change in flagella helicity) of flagella, and their arrangement at the body surface strongly influence the run-and-tumble behavior. Mesoscale hydrodynamics simulations with the developed model help us better understand physical mechanisms which govern *E. coli* dynamics, yielding the run-and-tumble behavior that compares well with experimental observations. This model can further be used to explore the behavior of *E. coli* and other peritrichous bacteria in more complex realistic environments.

## 1. Introduction

There exists a huge variety of microorganisms, which are part of nearly any environment, including air, water, soil, and biological organisms. Microorganisms generally have micrometer dimensions, and many of them possess some type of motility. Motile microorganisms are often referred to as microswimmers, which propel themselves to explore the environment and search for nutrition. In a fluid environment, propulsion of biological microswimmers is often facilitated by external appendages attached to the body such as flagella and cilia ^1–3^. Common examples are sperm cells with a beating flagellum ^4^, *E. coli* bacteria propelled by several rotating flagella ^5,6^, and paramecia whose body is covered by a large number of hair-like active cilia ^7^. The scientific interest in understanding the motile behavior of microswimmers ranges from fundamental characterization of physical mechanisms of propulsion ^1–3^ to the construction of microrobotic systems mimicking various aspects of biological microswimmers ^8,9^.

An interesting biological microswimmer is the bacterium *E. coli*, which employs multiple helical flagella for the propulsion ^5^. A typical *E. coli* has a sphero-cylinder-like body with a length of 2.5 *±* 0.6 *µm* and a diameter of 0.88 *±* 0.09 *µm* ^10^, and typically between 2 and 7 helical flagella with a length of 8.3 *±* 2.0 *µm* ^11^ attached to the body. Each flagellum is driven by a reversible rotary motor placed within the base membrane ^5,6^. When all flagella rotate in the same direction, they form a single bundle, which facilitates an efficient forward propulsion. A wild-type *E. coli* bacterium has a swimming speed of 29 *±* 6 *µm/s* in an aqueous environment ^10^. The synchronization between flagella and the formation of a tight bundle is facilitated by hydrodynamic interactions between flagella, as shown in several theoretical studies ^12–15^ and experiments ^16,17^. Flagella elasticity is also important for bundle formation, since it does not occur for rigid helices ^14,15^ .

To explore the environment and change the swimming direction, *E. coli* bacteria employ a so-called ‘run-and-tumble’ motion (see Fig. 1), which resembles a random walk ^5,23^. During a run phase, all left-handed flagella rotate in the anti-clockwise direction, and form a tight bundle that serves as a single helical propeller. Therefore, the run phase leads to a directed motion. In order to turn, *E. coli* bacteria ‘tumble’, where one or more flagella switch the rotation direction to clockwise and leave the bundle, facilitating a change in the swimming direction ^24,25^. As a result, *E. coli* can navigate through the environment using an interchangeable sequence of longer directional runs and shorter tumble phases ^24,26^. Note that in the presence of chemical gradients, the seemingly random run-and-tumble behavior of *E. coli* may change with a substantial reduction in the tumbling frequency ^27,28^.

**Figure 1.**
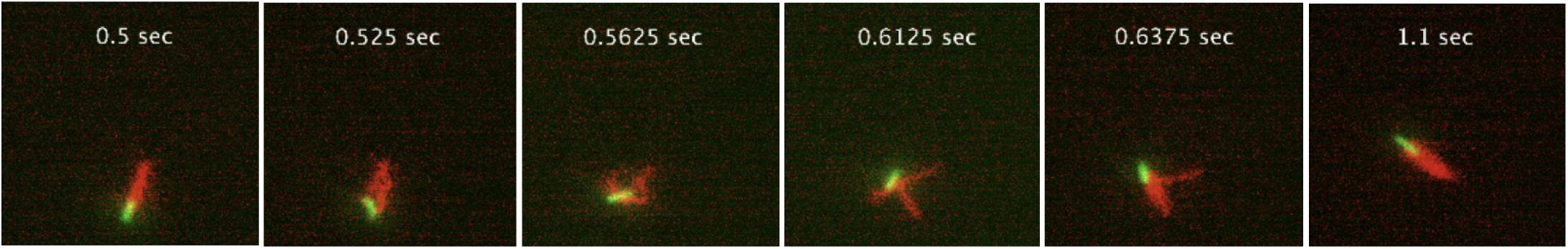
Experimental observation of a typical run-and-tumble dynamics of *E. coli*. Bacterium body is fluorescently labelled in green, while flagella are labelled in red. The run stage is when all flagella are in a single tight bundle, while the tumble stage is when one or several flagella leave the bundle. See also Movie S1.

Even though the qualitative description of *E. coli* tumbling is fairly simple, its physical realization is more complex. Experimental observations show that the flagella of *E. coli* can have several different polymorphic forms, depending on their state during the run and tumble phases ^10,24^ . During the run phase, all flagella have a shape referred to as a ’normal state’, which is a left-handed helix with a pitch of ∼ 2.5 *µm* and a diameter of ∼ 0.5 *µm*. However, upon the start of the tumble phase, one or two of the flagella change their helicity from left-handed to right-handed with different helical properties (e.g., pitch and diameter), in addition to the alteration of rotation in the clockwise direction ^24,25^; this is called a polymorphic transformation. At the end of the tumble phase, these reverted flagella change their rotation again from clockwise to anti-clockwise, thereby restoring the original state.

Another interesting aspect is the flexural rigidity of the flagellar hook, the short part of a flagellum that connects it directly to the motor. The hook plays an important role in flagella bundling and unbundling, and thus, contributes to the control of the swimming behavior ^29–31^. On the one hand, both theoretical and experimental studies ^11,32,33^ show that the hook must be sufficiently flexible to allow multi-flagellated bacteria to form a flagellar bundle. On the other hand, the hook must also be stiff enough to withstand forces exerted by the motor, and aid clockwise-rotating flagella in their separation from the bundle during tumble events. Mechanical properties of the hook have been measured, yielding a torsional rigidity of ∼ 10^7^ *Nm*^−2^ and a bending rigidity of ∼ 10^−29^ *Nm*^2 34^. Other investigations ^30,35,36^ report changes in the bending rigidity of the hook, depending on the load applied by the motor. For a steady swimming *E. coli*, a bending rigidity of 2.2 *×* 10^−25^ *Nm*^2^ was measured for the hook, while for a resting bacterium the value of 3.6 *×* 10^−26^ *Nm*^2^ was reported ^30^. Also, an increase in the hook bending rigidity from 5 *×* 10^−26^ *Nm*^2^ to 3 *×* 10^−24^ *Nm*^2^ was suggested with an increase in the motor torque ^35^. A further study reports the hook to be stiffer when the motor rotates clockwise in comparison to the anti-clockwise rotation under similar motor torques ^36^.

Since the first observations of the run-and-tumble behavior of wild-type *E. coli* ^5^, a number of experiments have been performed to quantify bacteria motion. The tumble angle, defined as a change in the swimming direction before and after a tumble event was estimated to be in the range 62^°^ − 68^°^ for a tumble time of 0.2*s* in Ref. ^5^, while tumble angles of 57 *±* 37^°^ for a tumble time of 0.19 *s* have been reported in Ref. ^37^. Furthermore, measurements of anti-clockwise/clockwise time series of a motor for wild-type *E. coli* estimate the time of clockwise rotation to be ∼ 0.38 *s* ^38^. Note that all these measurements may not have consistent definitions for the tumble angles and times.

To better understand experimental observations and the underlying mechanism, several simulation studies with detailed *E. coli* models have recently been performed ^19,39–41^. In Refs. ^39,40^, only run dynamics of *E. coli* for different bacterium properties and near surfaces was investigated. The run-and-tumble behavior was studied in Refs. ^19,41^, reporting an increasing tumble angle from 10^°^ to 80^°^ for an increasing tumble time. However, numerical discretization of *E. coli* body and flagella was quite coarse. Furthermore, the question of how different bacteria characteristics, such as body geometry, flagella and hook bending rigidity, and the polymorphic transformation, determine the run-and-tumble behavior, has not been directly considered.

In our work, we establish a detailed model of *E. coli* and investigate its run-and-tumble behavior in comparison to experimental observations. We show that the body geometry (sphero-cylinder-like or spheroidal) strongly affects the balance between rotational frequencies of the body and flagellar bundle, which may significantly limit successful tumbling of *E. coli*. Furthermore, both the polymorphic transformation and the stiffening of the hook of clockwise-rotating flagella are found to play important roles in an efficient change of the swimming direction. In particular, they govern the efficient separation of clockwise-rotating flagella from the bundle. Our model helps to better understand the importance of different *E. coli* characteristics for the run-and-tumble behavior, which is qualitatively consistent with the corresponding experimental observations. In the experiments, a novel two-color 3D Lagrangian tracking method ^42^ is employed for the observation of flagella configurations and dynamics of E. coli bacteria under free-swimming conditions in the bulk. Our computational model can further be used to investigate the behavior of *E. coli* and other peritrichous bacteria in complex environments, such as near surfaces and in complex geometries.

## 2 Methods and models

### 2.1 *E. coli* model

Our *E. coli* model is composed of two parts: (i) a rounded sphero-cylinder-like cell body and (ii) *N*_*f lag*_ left-handed helical flagella (see Fig. 2). The surface of the cell body, aligned with the *x* axis, is described by

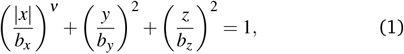

where *ν* = 8.5 for a sphero-cylinder-like shape, *b*_*x*_ = 1.5 *µm* is the major half-axis, and *b*_*y*_ = *b*_*z*_ = 0.5 *µm* is the minor half-axis, following the aspect ratio of *b*_*x*_*/b*_*y*_ ≈ 2 − 3 for wild *E. coli* ^10^. The body surface is represented by *N*_*v*_ = 1278 point particles that form an elastic triangulated network through bonds with a non-linear potential ^43,44^

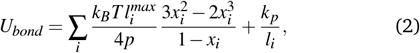

where 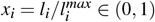, *l*_*i*_ is the length of bond 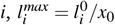is the maximum extension of spring *i, l*_*i*_ is the spring length from an initial triangulation of the body, and *x*_0_ = 0.45 is a constant for all springs. Furthermore, *p* is the persistence length, *k*_*p*_ is the repulsive force coefficient, and *k*_*B*_*T* is the thermal energy unit defined by the temperature *T* of the simulated system.

**Figure 2.**
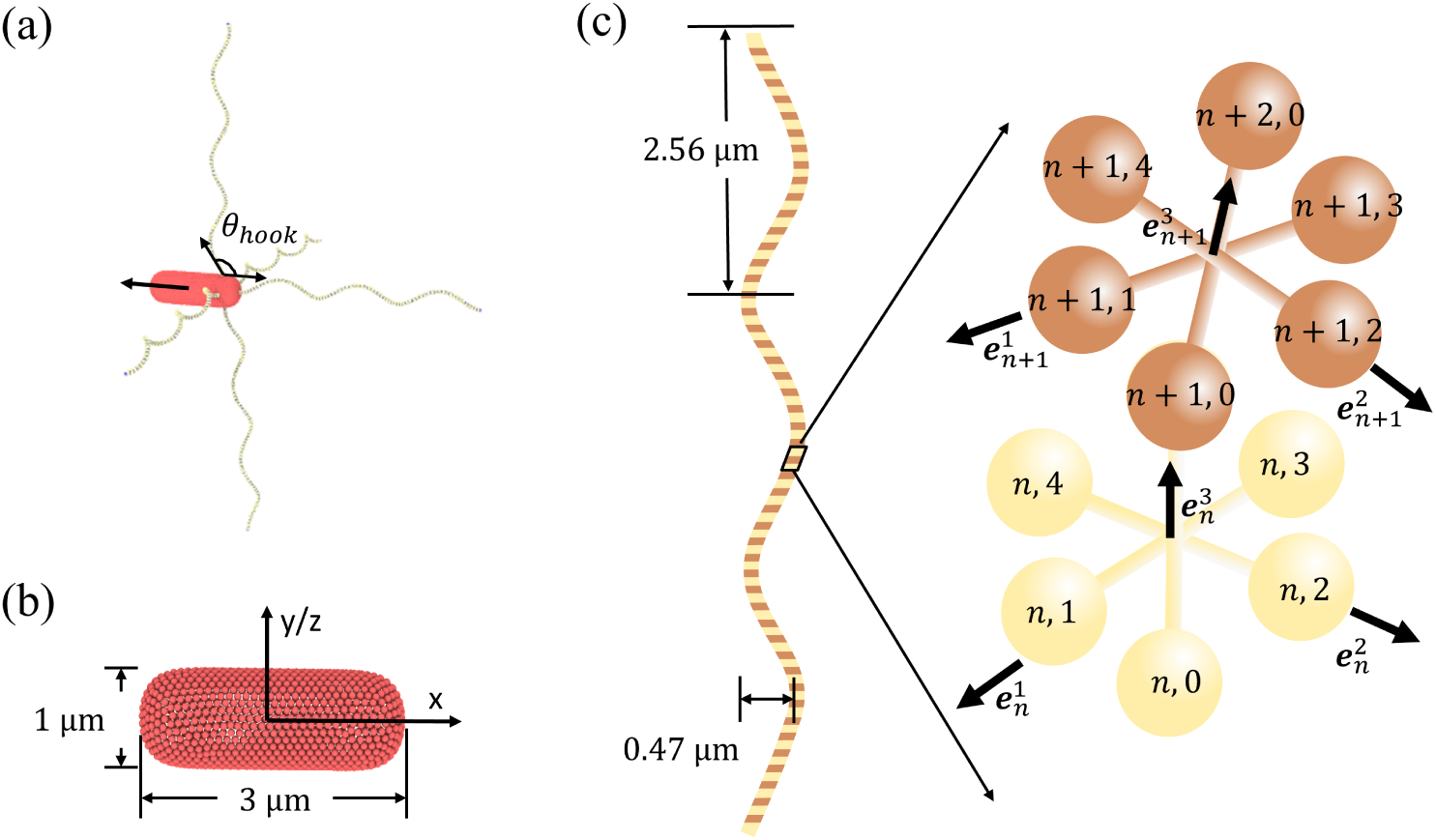
(a) *E. coli* model that consists of a sphero-cylinder-like cell body and *N*_*flag*_ = 5 left-handed helical flagella. The black arrow indicates the orientation vector of the body that is aligned with its major axis. The hook angle *θ*_*hook*_ is labelled and defined as the angle between the first few sections of a flagellum and the body surface. (b) Sphero-cylinder-like cell body described by Eq. (1) has a length of 3 *µm* and a diameter of 1 *µm*. It is represented by a collection of 1278 particles, forming a triangulated spring network on its surface. (c) Model of a left-handed flagellum with three helical turns made of *N*_*s*_ = 76 segments. It has a diameter of 0.47 *µm* and a pitch length of 2.56 *µm*. The flagellum model is adopted from Ref. ^39^.

In addition to the shear elasticity imposed by *U*_*bond*_ , the body shape is further stabilized by a curvature elasticity implemented through the Helfrich bending energy ^45^ discretized as ^46,47^

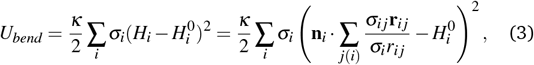

where *κ* is the membrane bending rigidity, *σ*_*i*_ is the area of particle *i* in a dual lattice, 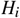 is the mean curvature at vertex *i*, and 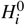 is the local spontaneous curvature. *σ*_*i*_ = ∑ _*j*(*i*)_ *σ*_*i j*_*r*_*i j*_*/*4, where *j*(*i*) spans all vertices linked to vertex *i, σ*_*i j*_ = *r*_*i j*_ (cot *θ*_1_ + cot *θ*_2_)*/*2 is the length of the bond in the dual lattice with *θ*_1_ and *θ*_2_ being the angles at the two vertices opposite to the edge *i j* in the dihedral. Furthermore, **r**_*i j*_ = **r**_*i*_ − **r** _*j*_ , *r*_*i j*_ = |**r**_*i j*_| , and **n**_*i*_ is the unit normal at the vertex *i*. Note that the spontaneous curvature 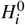 is prescribed locally after the triangulation of the body surface.

Conservation of the area and volume of the body is controlled by the potential ^43,44^

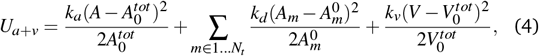

where *k*_*a*_, *k*_*d*_ and *k*_*v*_ are the coefficients of global area, local area, and volume conservation constraints. *A* and *V* are the instantaneous area and volume of the body surface,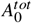 and 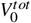 are the targeted global area and global volume which are set to those of the selected sphero-cylinder-like shape. *A*_*m*_ is the area of the *m* − *th* triangle (or face), while 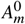 is its targeted value. *N*_*t*_ is the number of triangles within the triangulated surface. Note that the model described by the potentials above has been used to represent deformable and nearly rigid particles of various shapes ^43,48,49^.

The flagellum model is borrowed from Ref. ^39^. Figure 2(c) illustrates a left-handed helical flagellum comprised of *N*_*s*_ = 76 segments with 381 particles in total. Each segment consists of six particles arranged in a regular octahedral structure, stabilized by 12 springs with an equilibrium length of 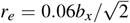. Opposite particles within the octahedral arrangement are connected by 3 diagonal springs with a preferred bond length of *r*_*d*_ = 0.06*b*_*x*_. The octahedral construction of a flagellum allows a straightforward imposition of the inherent helical twist, and facilitates a better coupling of the flagellum to the particle-based fluid.

The flagellar backbone is determined by the bonds 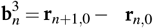 , where *n* = 1 … *N*. The backbone bonds, along with 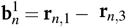 and 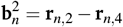, establish the corresponding orthonormal triads 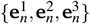, where 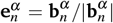 with *α* ∈ {1 2 3}. Here, **r**_*n*,0_ represent positions of the backbone particles, while **r**_*n*,*k*_ with *k* ∈ {1, 2, 3, 4} correspond to positions of the auxiliary particles lying in the plane perpendicular to the backbone vector 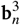.

To capture the local twist and bending of the flagellum, transformation of one triad 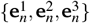 to the next 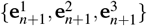 along the backbone is performed in two steps: (i) first,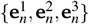 is rotated around 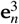 by a twist angle φ_*n n*_, and (ii) second, the twisted triad is rotated by a bending angle *θ*_*n*_ around the normal 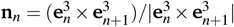 to the plane defined by the backbone bonds 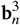 and 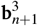. Following these transformations, an elastic deformation energy of the helical flagellum is defined as

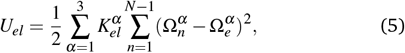

where 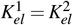 determine the bending rigidity and 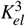 the twisting rigidity. 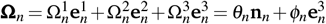 is the strain vector. The equilibrium structure of the flagellum is defined by the 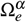 parameters in Eq. (5), which determine the helix radius and pitch length. Note that the handedness of the flagellum can simply be changed by altering the sign of 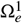 and 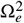.

Each flagellum is attached to the body by making the first particle **r**_1,0_ of the flagellum backbone to be part of the body discretization. To impose anchoring orientation of the flagellum with respect to the body, an angle potential

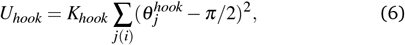

is employed, where *K*_*hook*_ is the potential strength, *i* is the anchoring particle at the body such that **r**_*i*_ = **r**_1,0_, *j*(*i*) spans all body vertices linked to vertex *i*, and 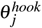 is the angle between vectors **r** _*j*_ − **r**_*i*_ and **r**_2,0_ − **r**_*i*_. The preferred angle of anchoring orientation is chosen to be *π/*2, imposing a perpendicular orientation of the flagellum to the body. The potential *U*_*hook*_ mimics the rigidity of a flagellum hook, which is an initial short part (50 − 60 nm in length) of the flagellum that is generally softer than the rest of the flagellum ^30,50^. Thus, *K*_*hook*_ represents the bending rigidity of the physical hook, and will be referred to as a hook rigidity below.

The action of a motor imposing flagellum rotation is implemented by a torque **T**_*m*_ that is aligned with 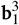 , and exerted onto the particles 1, *k* ∈ ({*k* 1, 2, 3, 4 }) of the first segment of the flagellum. To satisfy torque-free and force-free conditions for the swimmers, a counter torque − **T**_*m*_ is imposed onto the body particles *j*(*i*). Finally, excluded-volume interactions between different flagella, and between flagella and the body surface are implemented using the repulsive part of the Lennard-Jones (LJ) potential as

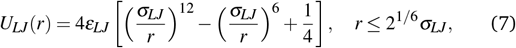

where *ε*_*LJ*_ sets the strength of the potential, *σ*_*LJ*_ is the characteristic repulsion length, and *r* is the distance between two interacting particles.

### 2.2 Modeling fluid flow

Fluid flow is modeled by the dissipative particle dynamics (DPD) method, a mesoscopic hydrodynamics simulation technique ^51,52^. The DPD fluid consists of a collection of particles, each representing a small fluid volume. DPD particles *i* and *j* interact through three pair-wise forces (conservative, dissipative, and random)

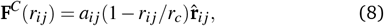

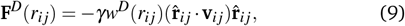

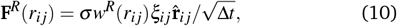

where *a, γ*, and *σ* determine the force strengths, **r**_*i j*_ = **r**_*i*_ − **r** _*j*_ is the distance vector, *r*_*i j*_ = |*r*_*i j*_| , 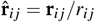, and **v**_*i j*_ = **v**_*i*_ − **v** _*j*_ is the velocity difference. Δ*t* is the time step, and *ξ*_*i j*_ = *ξ* _*ji*_ is the symmetric Gaussian random variable with zero mean and unit variance. The dissipative **F**^*D*^ and random **F**^*R*^ forces define a thermostat and are related to each other through the fluctuation-dissipation theorem ^52^ as

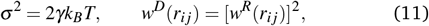

with the weight function *w*^*R*^(*r*_*i j*_ )

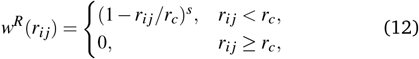

where the exponent *s* controls the decay of *w*^*R*^(*r*_*i j*_ ) as a function of inter-particle distance. All forces vanish beyond the cutoff radius *r*_*c*_.

The repulsive force **F**^*C*^ controls fluid compressibility, while the dissipative force **F**^*D*^ reduces the velocity difference between two neighboring particles, controlling fluid viscosity. The positions and velocities of DPD particles are updated using the Newton’s second law

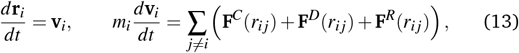

where *m*_*i*_ is the mass of particle *i*. The time integration in our simulations is performed using the velocity-Verlet algorithm ^53^.

### 2.3 Simulation setup and parameters

We define a length scale as the length of the major half-axis *b*_*x*_ (*b*_*x*_ = 9 in simulation units), a time scale *τ* (*τ* = 132 in simulation units) as the period of bundle rotation, and an energy scale as *k*_*B*_*T* (*k*_*B*_*T* = 1 in all simulations). Simulations are performed in a domain with dimensions 8.33*b*_*x*_ *×* 11.56*b*_*x*_ *×* 11.56*b*_*x*_, where periodic boundary conditions are imposed in all three directions. Each simulation contains a single bacterium and 2.4 *×* 10^6^ fluid particles with a number density of 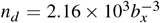. DPD parameters for interactions between fluid particles are *a* = 540*k*_*B*_*T/b*_*x*_, 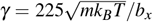, *s* = 0.15, and *r*_*c*_ = 0.11*b*_*x*_ (*m* = 1 in all simulations). These parameters result in a dynamic viscosity of 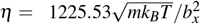. The simulations are run for a total duration of approximately 113.64*τ* with a time step Δ*t* = 2.27 *×* 10 ^−5^*τ*.

In simulations, we set 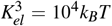 , which is large enough to prevent significant twisting of flagella during rotation. 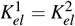 are set in the range between 2 *×* 10^3^*K*^*B*^*T* and 5 *×* 10^4^*K*_*B*_*T* , corresponding to bending stiffnesses 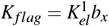 between 1.7 *×* 10^−23^ *Nm*^2^ and 4.2 *×* 10^−22^ *Nm*^2^. These values are within the range of experimentally measured bending stiffnesses 10^−24^ − 10^−21^*Nm*^2 39^. Furthermore, 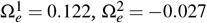, and 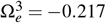, defining a left-handed helix in equilibrium with a radius of 0.23*µm* and a pitch length of 2.56*µm* ^10^. Each flagellum consists of *N*_*s*_ = 76 segments, resulting in three helical turns. A basic *E. coli* model has five flagella, with one attached at one body end and four attached symmetrically to the side closer to the end [see Fig. 3(a)]. All flagella are actuated by a constant torque *T*_*m*_ applied to the first segment of flagellum structure. Note that for real *E. coli*, the motor torque may depend on the hydrodynamic load ^54,55^, and is not a control parameter directly, but here, to keep numerical simulations simple, we chose to keep *T*_*m*_ constant.

**Figure 3.**
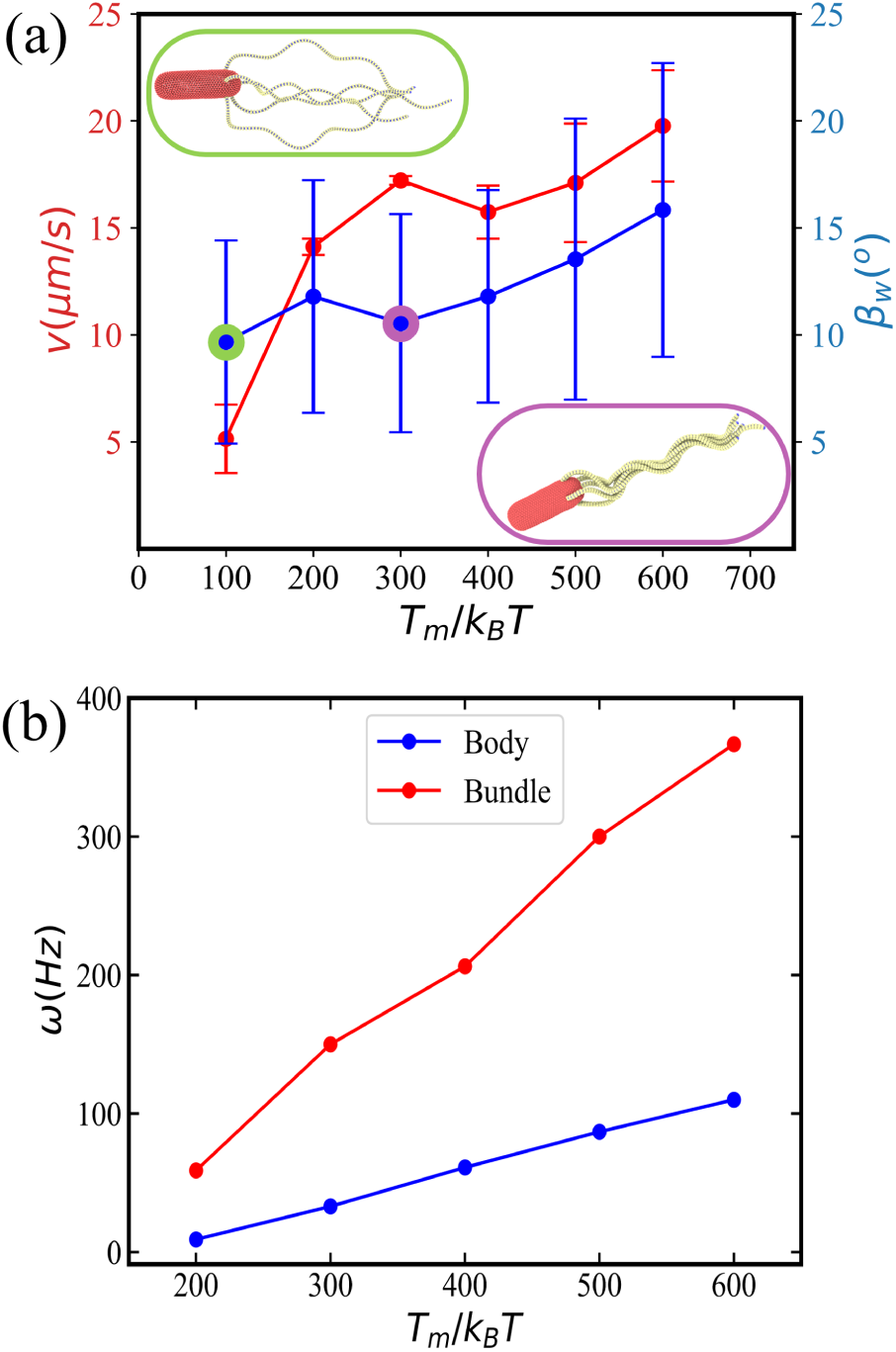
Five flagella *E. coli* model for different values of torque. (a) Average swimming speed *v* and wobbling angle *β*_*w*_ as a function of applied torque *T*_*m*_. The swimming speed is computed from a fixed-time displacement and the wobbling angle is defined as the angle between the orientation vector of the body and the axis of flagellar bundle during forward swimming (i.e., run phase). Insets show snapshots of *E. coli* for the applied torques *T*_*m*_ = 100*k*_*B*_*T* (top) and *T*_*m*_ = 300*k*_*B*_*T* (bottom) with a poorly formed and tight flagellar bundle, respectively (see also MoviesS2 and S3). (b) Rotation frequencies of the body and the bundle as a function of torque.

The body of *E. coli* is discretized by *N*_*v*_ = 1278 particles. The corresponding surface has a shear modulus of *µ*_0_ = 8.1 *×* 10^4^*k*_*B*_*T/b*^2^ and the bending modulus *κ* = 100*k*_*B*_*T* . The area and volume constraints assume 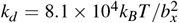,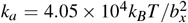, and 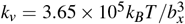 , with the body area 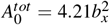 and the body volume 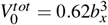. Excluded-volume interactions between the body and flagella or between different flagella are implemented through the LJ potential with parameters *ε*_*LJ*_ = *k*_*B*_*T* and *σ*_*LJ*_ = 0.07*b*_*x*_. Coupling between the *E. coli* model and fluid flow is imposed through DPD interactions with the parameters *a* = 0, 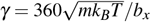 , *s* = 0.1, and *r*_*c*_ = 0.09*b*_*x*_. A characteristic Reynolds number *Re* = *b*_*x*_*mn*_*d*_*v/η*≈ 0.01 is small enough to neglect inertial effects. Here,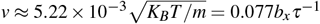 is the swimming speed of *E. coli*.

Simulations are carried out in two steps. First, *E. coli* with initially unbundled flagella (all rotating anti-clockwise) is let to swim and form a tight bundle due to hydrodynamic attraction between them ^12,13,16,17^. Subsequently, we alternate between the run phase with all flagella rotating anticlockwise and the tumble phase when one or two pre-selected flagella switch to the clockwise rotation through altering the torque direction. The run phase takes about 18.9*τ* and the tumble phase approximately 12.6*τ*, so that the duration of each simulation corresponds to approximately 3 tumble events. Note that the time scale *τ* depends on the applied torque *T*_*m*_ to each flagellum. In most simulations, the pre-selected flagella are also subject to polymorphic transformation, where the flagellum helicity is changed from left-handed to right-handed. We employ a simplified version of polymorphic transformation with only two states, left-handed and right-handed helices with the same helix radius and pitch. The polymorphic transformation is implemented such that the parameters 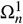 and 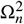 are changed from their original values (representing a left-handed helix) to the same magnitudes with the opposite sign (representing a right-handed helix) in a linear fashion during time 0.76*τ*. When the tumble phase is completed, the pre-selected flagella are again subject to anti-clockwise rotation and the reverse polymorphic transformation (from a right-handed to a left-handed helix), leading to the formation of the flagellar bundle.

To compare simulation results with experimental observations, we assume the body length of 3*µm* and the body diameter of 1*µm* ^10^. The period of bundle rotation during *E. coli* run is *τ* = 6.7 *×* 10^−3^*s* with a rotation frequency of 150 Hz ^10^, which allows us to relate simulation and physical time scales. This means that each simulation corresponds to a duration of 0.76*s*, with the run and tumble times of approximately 0.13 s and 0.08 s, respectively. Note that the run time of 0.13 s is shorter than typical *E. coli* run times of about 1 s, which has been selected to reduce computational cost, since we primarily focus on *E. coli* tumbling. However, the time of 0.13 s is long enough to form a tight flagellar bundle after a tumbling event. Ambient conditions correspond to a temperature of *T* = 20^*o*^ C with the fluid viscosity of *η* = 10^−3^*Pa · s*.Table 1 compares the properties of simulated and real *E. coli*.

**Table 1.** *E. coli* parameters used in simulations and measured experimentally with the corresponding references.

| Property | Simulation | Experiment | Reference |
| --- | --- | --- | --- |
| Body length ($\mu m$) | 3.0 | $2.5 \pm 0.6$ | 10 |
| Body width ($\mu m$) | 1.0 | $0.9 \pm 0.6$ | 10 |
| Flag. contour length ($\mu m$) | 8.9 | $8.3 \pm 2.0$ | 56 |
| Flag. pitch length ($\mu m$) | 2.6 | 2.2 | 56 |
| Flag. helix radius ($\mu m$) | 0.2 | 0.2 | 56 |
| Flag. bending rigidity ($Nm^2$) | $10^{-22}$ | $10^{-24} - 10^{-21}$ | 39 |
| Mean run time (s) | 0.13 | 1.0 | 5,10 |
| Mean tumble time (s) | 0.08 | $0.14 \pm 0.08$ | 24 |

### 2.4 Experimental methods and protocols

E. coli strains used in the experiments are mutant strains AD62 and AD63. Both were genetically modified to express green fluorescent protein (GFP) in their body, while the flagella are specifically labelled with a fluorescent protein Alexa 647. This two-color technique prevents signal overlap and enables a distinct simultaneous observation of both the bacterial body (green) and flagella (red) via our original 3D Lagrangian tracking technique ^42^. From the perspective of run-and-tumble motility, AD62 corresponds to a standard wild-type E.coli. It was used to visualize the tumbling dynamics. AD63 is a Δ*CheY* mutant of AD62 that does not tumble (“smooth-runner”). It was used to study the relation between swimming velocity and the number of flagella. For this last experiment, each bacterium was tracked individually at a surface using a single color (GFP) for a time long enough (around 50 s) to obtain a precise measurement of the swimming velocity. After that, a UV-light was shined directly on the bacterium to obtain enough photo damage (after about 100 s), so that it stops the motion and triggers a full debundling of the flagella. The second color excitation is then triggered to visualize and count the flagella individually.

The Lagrangian tracking device comprises of two super-imposed stages mounted on an inverted epifluorescence microscope (Zeiss Observer Z1, equipped with a C-Apochromat 63 *×* 1.2*W* objective). The horizontal (*x, y*) position is controlled mechanically, and the *z* − position is adjusted using a piezo-electric mover. A targeted fluorescent bacterial body is visualized within a “trapping area” using a CCD camera. To synchronize image acquisition with stage movements, the stages and camera are triggered by a National Instruments TTL trigger module. The acquired images are transferred to a Labview program, which processes the data. The program records the current x, y, and z positions and directs the mechanical and piezoelectric stages to adjust accordingly, ensuring that the particle remains within the “trapping area” and in focus ^57^.

Furthermore, to simultaneously image both the fluorescent body and the flagella, a two-color LED light source (Zeiss Colibri 7) and a dichroic image splitter (Hamamatsu) are utilized to project two monochrome images onto separate regions of the camera chip. The microscope stage movement is computer-controlled to maintain the selected bacterium in focus, with images (1024 *×* 1024 pixels) captured at 80 frames per second using a Hamamatsu ORCA-Flash 4.0*C*11440 camera. The green and red images are subse-quently superimposed to generate a movie depicting the tracked bacterium and its flagellar bundle (see Movie S1). Due to photo-bleaching, flagella imaging is limited to approximately one minute, but the timing of the second color channel’s application can be controlled to visualize flagella dynamics as needed ^42^.

The bacterial suspension must be dilute enough to avoid frequent bacterial collisions, and is prepared according to the following protocol. Bacteria are inoculated in 10*mL* of Luria Broth (LB) containing ampicillin at a concentration of 100*µg/mL* and incubated overnight at 30^°^*C*. Subsequently, 100*µL* of this culture is transferred into 10*mL* of Tryptone Broth (TB) and incubated for several hours until the bacteria reach the early stationary phase. The bacteria are then harvested by centrifugation, and the supernatant is discarded. The pellet is resuspended in 1*mL* of Berg Motility Buffer (BMB: 6.2*mMK*_2_*HPO*_4_, 3.8*mMM*_2_*PO*_4_, 67*mMNaCl*, and 0.1*mMEDTA*) containing 10*µL* of Alexa Fluor 647 red dye (prepared as a 5*mg/mL* stock solution in Dimethylsulfoxide (DMSO)). The suspension is gently shaken for 2 hours. Following this 2-hour dyeing process on the bacterial flagella, the cells are washed again by centrifugation. The supernatant is then removed, and the final pellet is resuspended in a specific volume of BMB solution containing PVP (with 0.08g/mL L-serine) to maintain the activity of bacteria and to inhibit bacterial division.

## 3. Results

### 3.1 Run phase

#### 3.1.1 Flagellar bundle formation

During the run phase, when all flagella rotate anti-clockwise, they form a tight bundle due to the rotational flow field between them and body counter-rotation ^12,13,15–17^. However, hydrodynamic interaction forces are relatively weak and the formation of the bundle may not take place for small torques applied. Insets in Fig. 3(a) illustrate two snapshots of *E. coli* with *T*_*m*_ = 300*k*_*B*_*T* and *T*_*m*_ = 100*k*_*B*_*T* , where a tight bundle is formed or does not form, respectively (see also Movies S2 and S3). Our simulations show that torques *T*_*m*_ ≳ 200*k*_*B*_*T* lead to the successful formation of a tight bundle. Note that the tightness of the bundle also depends on flagellar bending rigidity, so that a sufficiently small rigidity is necessary for its formation ^12^. Furthermore, the hook rigidity plays an important role in bundle formation, such that stiff hooks may prevent the formation of a bundle due to a preferred perpendicular orientation of flagella with respect to the body surface. This is consistent with experimental measurements that the hook is significantly softer than the rest of flagella ^11,32,33^.

#### 3.1.2 *E. coli* propulsion as a function of applied torque

To quantify the run dynamics, we measure *E. coli* swimming speed *v* and wobbling angle *β*_*w*_. The swimming speed is computed from a fixed-time displacement as *v* = |**r**(*t*_0_ + Δ*t*) − **r**(*t*_0_) |*/*Δ*t*, where **r**(*t*) is the center of mass of *E. coli* at time *t*, and Δ*t* is a fixed time difference during which the bacterium does not change its swimming direction. We use Δ*t* = 22.8*τ* (0.15 s) for simulations with the bacterium remaining in the run phase (i.e., no tumbling). The wobbling angle *β*_*w*_ corresponds to the angle between the body axis and a vector defined by the orientation of the bundle, and the angle ranges from 0 to 90^°^. Figure 3(a) shows the relationship of swimming speed and wobbling angle as a function of applied torque. The increase of speed with increasing torque is not linear. A large increase in *v* from 5 *µm/s* at *T*_*m*_ = 100*k*_*B*_*T* to 15 *µm/s* at *T*_*m*_ = 200*k*_*B*_*T* is due to the formation of a tight bundle at larger torques. For *T*_*m*_ *>* 200*k*_*B*_*T* , the increase in velocity is moderate and near linear as a function of torque. However, the wobbling angle also increases with increasing torque, which is expected to lead to an increased fluid resistance. Due to changes in *β*_*w*_, an increase in the swimming speed may not necessarily be linear as a function of applied torque.

Note that the swimming speed of 17.2 *µm/s* at *T*_*m*_ = 300*k*_*B*_*T* is smaller than the average *E. coli* speed of 29 ± 6 *µm/s* observed in experiments ^10^. This difference might be due to several reasons. First, a finite size of the simulation domain results in a reduction of swimming speed, since the modeled bacterium interacts hydrodynamically with itself through periodic boundary conditions. However, much larger simulation domains quickly become very expensive computationally, which would significantly limit our study of *E. coli* tumbling. Second, the thickness of simulated flagella is around 90 nm which is larger than in experiments (20 nm), resulting in a slightly reduced propulsion strength. Such high resolution is difficult to achieve in our simulations due to a high computational cost.

Figure 3(b) presents rotation frequencies *ω* of the body and the bundle as a function of applied torque. As expected, the both frequencies increase linearly with increasing *T*_*m*_. This also suggests that body wobbling does not significantly affect the rotation frequency of the body. Experimentally measured values are 23 ± 8 Hz for the body and 131 ± 31 Hz for the bundle ^10^, which agree well with *ω* values of 33 Hz and 150 Hz in simulations for *T*_*m*_ = 300*k*_*B*_*T* . Therefore, the *E. coli* model with five flagella and *T*_*m*_ = 300*k*_*B*_*T* is considered as a base model in further simulations.

#### 3.1.3 *E. coli* with different number of flagella

To test whether the number of flagella (typically between 2 and 7) affects dynamical characteristics of *E. coli*, we perform simulations for *E. coli* models with a different number of flagella. In all models, the arrangement of flagella is such that one flagellum is placed at the back along the long axis of the body, and the others are symmetrically placed at the selected circumference of the body surface. Figure 4(a) shows the swimming speed and the wobbling angle as a function of the number of flagella (see also Movies S4 and S5). The swimming speed remains nearly constant up to about five flagella and then slightly decreases for 6 and 7 flagella. A possible reason for the slight decrease in *v* is an increase in the wobbling angle with increasing number of flagella. Furthermore, our simulations indicate that the bundle becomes more spread with increasing number of flagella. Figure 4(b) presents rotational frequencies of the body and the bundle for different flagella numbers. The rotation of the body is nearly independent of the number of flagella, while the frequency of bundle rotation slightly increases.

**Figure 4.**
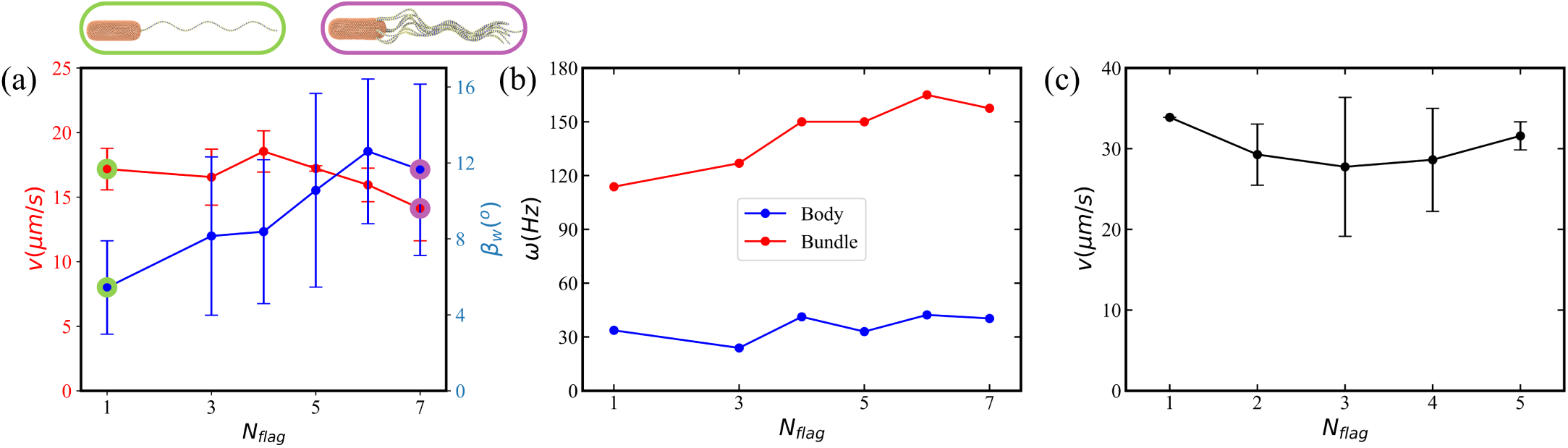
Dynamic properties of *E. coli* with different number of flagella (*N*_*flag*_ = 1, 3 − 7) during the run phase. *T*_*m*_ = 300*k*_*B*_*T* in all cases. (a) Swimming speed *v* and wobbling angle *β*_*w*_ for a simulated *E. coli* as a function of the number of flagella (see Movies S4 and S5 for *N*_*flag*_ = 1 and *N*_*flag*_ = 7). (b) Simulated rotation frequencies *ω* of the body and the bundle for different *N*_*flag*_. (c) Swimming speed of *E. coli* near a surface obtained from experiments.

For comparison, Fig. 4(c) shows the swimming speed of *E. coli* near a surface measured in experiments (see Section 2.4 for description). There is no obvious dependence of *v* on the number of flagella, which is consistent with simulations. A simple theoretical argument to support this observation is that the number of flagella only affects the thickness of the formed bundle. Since the propulsion strength would be expected to decrease weakly as a function of the bundle thickness, this effect might be very difficult to measure reliably. Furthermore, it is not clear whether the overall output power of *E. coli* increases with the number of flagella. In simulations, this is the case since each flagellum is driven by aconstant torque of *T*_*m*_ = 300*k*_*B*_*T* . Nevertheless, an increased out-put power with increasing number of flagella has nearly negligible effect on the swimming speed.

### 3.2 Tumble phase

Next, we study systematically the tumble phase, focusing on the importance of various bacterium characteristics for a successful change in the swimming direction. Here, one of the flagella rotates clockwise, in contrast to the bundle formed by other flagella rotating anti-clockwise.

#### 3.2.1 Effect of body shape

The shape of *E. coli* observed experimentally is generally sphero-cylinder-like (well described by Eq. (1) with *ν* = 8.5) with some variation in the body length. This slender shape somewhat maximizes the body volume for a fixed length and aspect ratio. For comparsion, we also consider a spheroidal body with the same dimensions defined by Eq. (1) with *ν* = 2, see Fig. 5(a). The *E. coli* model with the spheroidal body shape yields different rotation frequencies of the body and the bundle in the run phase ( ∼ 38 Hz and ∼ 75 Hz, respectively), compared to the model with the sphero-cylinder-like body. This is primarily due to different angles of flagella attachment and different fluid frictions on the body during rotational motion. Note that for the sphero-cylinder-like body, the side flagella have a preferred orientation perpendicular to the body axis in equilibrium [see Fig. 2(a)], while for the spheroidal body, the side flagella are partially aligned with the body axis. The ratio between bundle and body rotation frequencies significantly affects bacterium tumbling behavior. For the spheroidal body, this ratio is close to two, so that the body rotation is relatively fast, often resulting in wrapping of the body by the clockwise-rotating flagella, see Fig. 5(b) and Movie S6. Occurrence of the wrapping leads to a very limited change in the swimming direction. For the sphero-cylinder-like body, this ratio is close to five ^10^, such that the clockwise-rotating flagella are able to leave the bundle without significant wrapping of the body, resulting in an efficient change of swimming direction. The wrapping of clockwise-rotating flagella around the body for the spheroidal case can be prevented by reducing the applied torque; however, much smaller torques significantly affect the bundle formation and the propulsion strength of *E. coli*.

**Figure 5.**
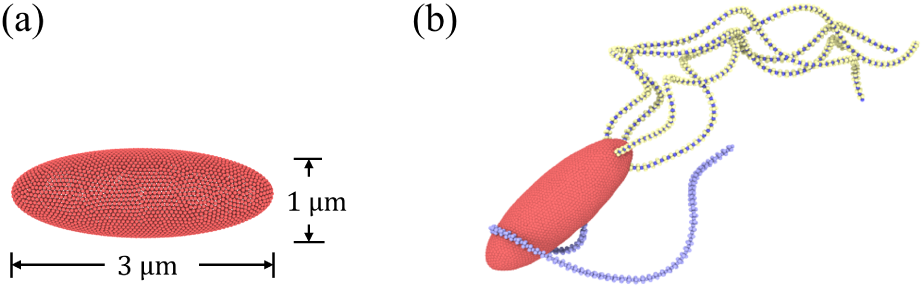
*E. coli* model with a spheroidal body. (a) Dimensions of the body with a spheroidal shape. (b) Illustrative snapshot of a bacterium during tumbling (see also Movie S6). Here, *N*_*flag*_ = 5, *K*_*flag*_ = 2.7 *×* 10^5^*k*_*B*_*Tb*_*x*_, *K*_*hook*_ = 500*k*_*B*_*T* , and *T*_*m*_ = 300*k*_*B*_*T* . The reverted flagellum does not undergo polymorphic transformation and hook stiffening.

#### 3.2.3 Straight initial section of flagella

Flagellum geometry plays an important role in bundling and unbundling processes during the run and tumble phases ^12,13,16,17^. We have tested the effect of a straight initial section of flagella (i.e., no helicity) near the attachment point to the body (not a hook). Figure 6 compares the run and tumble phases of *E. coli* models with a varying length *l*_*n*_ (from 0% to about 26% of the contour length) of the linear flagellum section. In all cases, the formation of a tight bundle during the run phase is successful, though the initial section of the bundle is somewhat loose for the cases with a non-zero length of an initial straight section. However, the tumbling behavior is significantly affected by the presence of a straight flagellum section, as can be seen from the snapshots in the bottom row in Fig. 6. Our simulations show that the clockwise-rotating flagellum has difficulties to leave the bundle for the initial straight sections of *l*_*n*_ = 10 − 20 segments (13 − 26% of the contour length). This result suggests that the larger the differences between the clockwise-rotating flagellum and the bundle, the easier for it to leave the bundle. For the case of no initial straight section, the mismatch of rotation is present for the whole contour length, facilitating the easier escape of the clockwise-rotating flagellum from the bundle.

**Figure 6.**
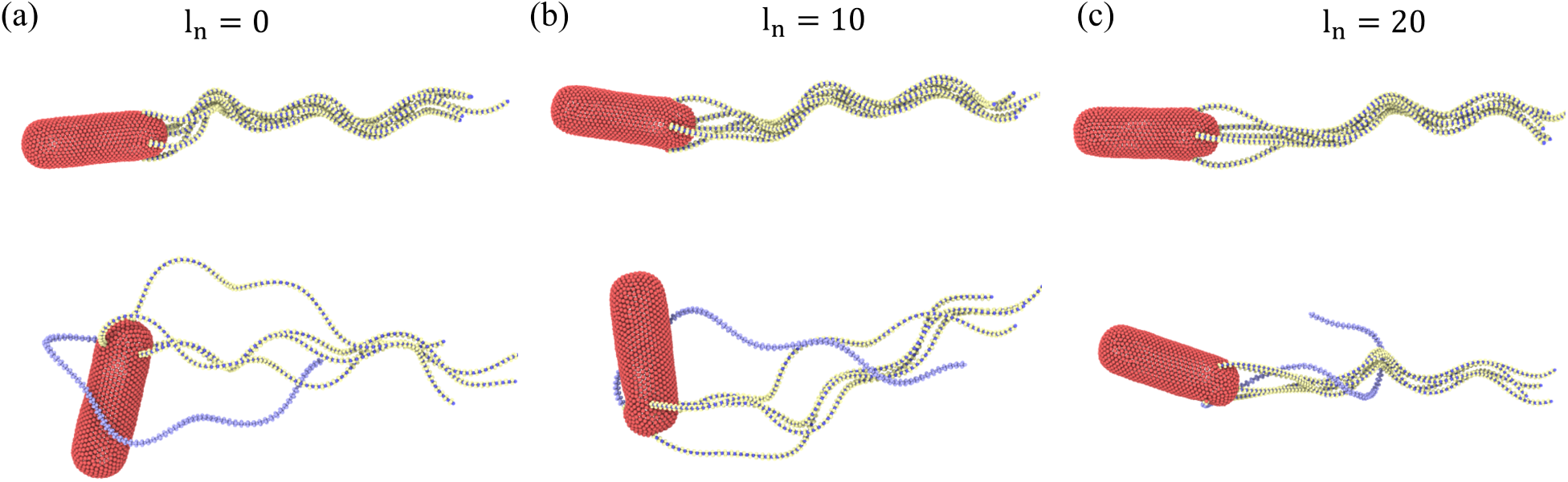
*E. coli* models with a varying length of the linear flagellum section at the attachment to the body. (a) No initial linear section with full three turn left-handed helices. (b) An initial linear section of *l*_*n*_ = 10 segments (13% of the contour length). (c) An initial linear section of *l*_*n*_ = 20 segments (26% of the contour length). Snapshots in the top row show the run phase, while the bottow row illustrates the tumble phase. In all models, *N*_*flag*_ = 5, *K*_*flag*_ = 2.7 *×* 10^5^*k*_*B*_*Tb*_*x*_, *K*_*hook*_ = 200*k*_*B*_*T* , and *T*_*m*_ = 300*k*_*B*_*T* . In all three cases, no polymorphic transformation and hook stiffening are incorporated.

#### 3.2.3 Polymorphic transformation

The snapshot of tumbling *E. coli* in Fig. 6(a) indicates that the bundle of anti-clockwise rotating flagella becomes loose when the clockwise-rotating flagellum leaves the bundle. This means that the clockwise-rotating flagellum significantly disturbs the bundle. This appears to be different in experimental observations where the bundle generally remains tight during the tumble phase.

Several studies ^10,24,25^ report that the clockwise-rotating flagella of a wild-type *E. coli* are subject to the polymorphic transformation, during which the flagella change their helicity from left-handed to right-handed. We implement this transformation by gradually changing the sign of original values of 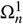 and 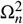 during a time of 0.76*τ*. An instant or very fast change of flagellar helicity is not stable in simulations. Figure 7 presents a snapshot of tumbling *E. coli* with polymorphic transformation, where the tightness of the bundle is not significantly affected by the leaving flagellum (see also Movie S7). At the end of the tumble phase, the reverted flagellum switches the rotation direction to anti-clockwise, performs the reverse polymorphic transformation (i.e., from the right-handed to left-handed helicity), and quickly joins the bundle.

**Figure 7.**
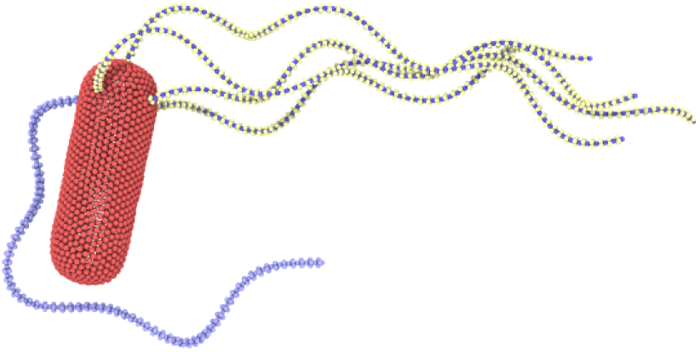
Tumble behavior of *E. coli* with polymorphic transformation of the reverted flagellum (see also Movie S7). Here, *N*_*flag*_ = 5, *K*_*flag*_ =2.7 *×* 10^5^*k*_*B*_*Tb*_*x*_, *K*_*hook*_ = 200*k*_*B*_*T* , and *T*_*m*_ = 300*k*_*B*_*T* . Hook stiffening is not incorporated.

The result that a polymorphic transformation of the clockwise-rotating flagella facilitates their efficient escape from the bundle without its significant disturbance can be understood as follows. A clockwise rotating flagellum that leaves the bundle without poly-morphic transformation exerts a propulsion force in the opposite-to-swimming direction, which competes with the propulsion force of the bundle. Since the propulsion forces from a single flagellum and a flagella bundle are comparable (supported by nearly independence of the swimming speed of *E. coli* on the number of flagella in Fig. 4), their counter-action in case of no polymorphic transformation of the clockwise-rotating flagellum leads to a significant slow-down of the bacterium, and thus, a partial disappearance of the bundle. However, in the case of polymorphic transformation, propulsion forces from this flagellum and the bundle act in a similar direction, so that the bacterium does not significantly lose its speed, maintaining a tight confirmation of the bundle. Furthermore, we find that the clockwise-rotating flagellum with polymorphic transformation does not attract the counter-rotating bundle hydrodynamically, further facilitating its efficient escape from the bundle.

#### 3.2.4 Stiffening of the hook during tumbling

Several recent studies ^35,36^ suggest that the hook of a reverted flagellum becomes stiffer during the tumble phase. Simulations with a fully flexible hook (i.e., *K*_*hook*_ = 0) lead to a poor tumbling of *E. coli*, because the initial section of the reverted flagellum is nearly aligned with the body surface. Even though the reverted flagellum separates from the bundle, it does not succeed to go sufficiently away from the bundle to significantly change the swimming direction. When the hook rigidity is increased, the situation drastically changes. Due to a preferred perpendicular orientation of the initial section of the reverted flagellum with respect to the body surface, its clockwise rotation pulls the flagellum out of the bundle. Furthermore, the hook rigidity drives the reverted flagellum toward a perpendicular-to-the-body orientation that is favorable for a proper tumbling. However, hook stiffness can also be not too large, as it would then prevent the formation of a tight bundle.

Figure 8 shows tumbling *E. coli* models for several hook rigidities. As expected, the larger the hook rigidity, the easier the reverted flagellum can leave the bundle. For *K*_*hook*_ = 100*k*_*B*_*T* in Fig. 8(a), orientation of the reverted flagellum is not well controlled, so that it partially wraps around the body. For *K*_*hook*_ = 200*k*_*B*_*T* in Fig. 8(b), orientation of the tumbling flagellum becomes better, but it still primarily remains near the bacterium body. Pronounced separation of the reverted flagellum is achieved for *K*_*hook*_ = 500*k*_*B*_*T* in Fig. 8(c); however, the bundle is relatively loose due to the large rigidity of flagella hooks. To incorporate hook stiffening due to different rotation directions into the model, we set *K*_*hook*_ = 500*k*_*B*_*T* for the reverted flagellum and *K*_*hook*_ = 100*k*_*B*_*T* for all other flagella. Figure 8(d) shows a tumbling event for this model, where a good separation of the reverted flagella with a minimum disturbance to the bundle is observed (see also Movie S8). In conclusion, stiffening of the hook of the reverted flagella facilitates *E. coli* tumbling.

**Figure 8.**
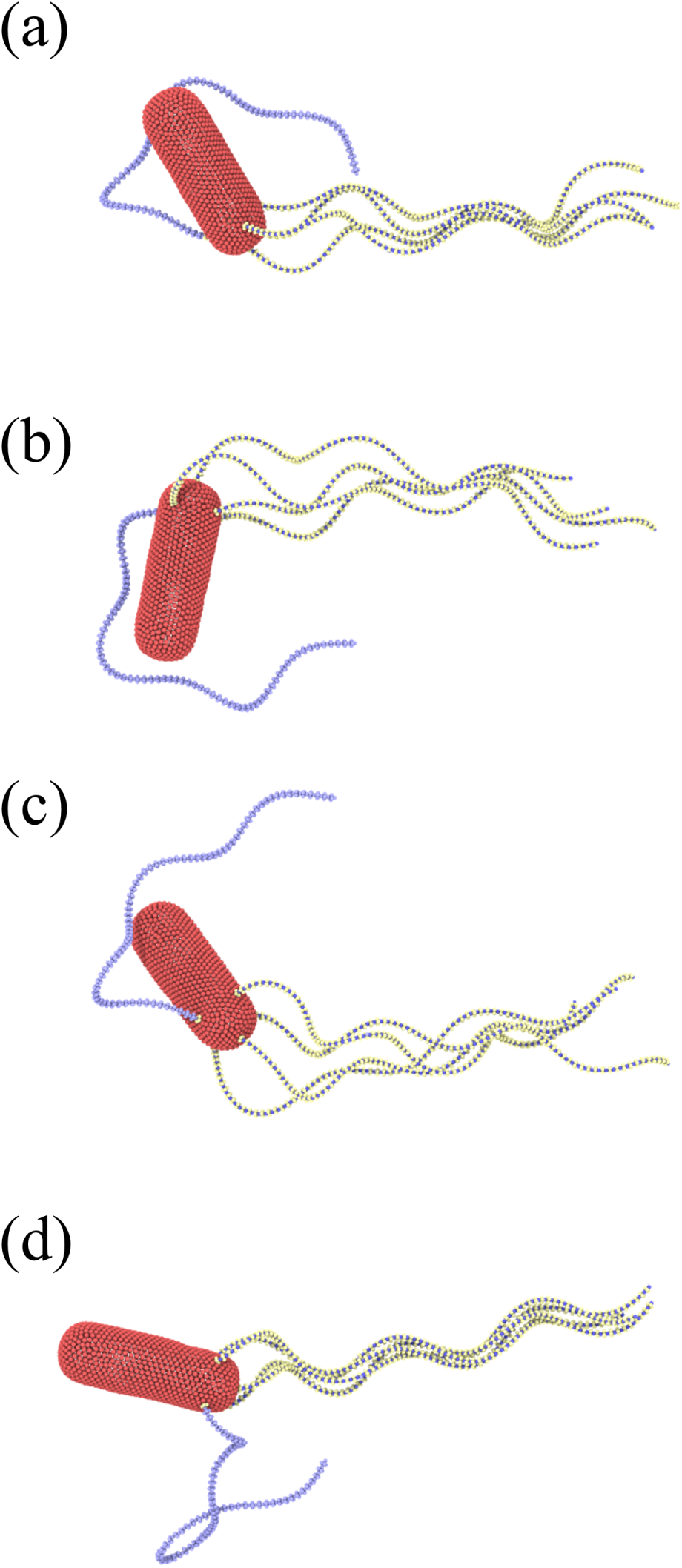
Tumble behavior of *E. coli* models with different hook rigidities.(a) All flagella have a hook rigidity of 100*k*_*B*_*T* . (b) All flagella possess *K*_*hook*_ = 200*k*_*B*_*T* . (c) All flagella have a hook rigidity of 500*k*_*B*_*T* . (d) A model where the reverted flagellum has *K*_*hook*_ = 500*k*_*B*_*T* , while the other flagella possess a hook rigidity of 100*k*_*B*_*T* (see also Movie S8). In all cases, the polymorphic transformation of the clockwise-rotating flagellum is implemented. Here, *N*_*flag*_ = 5, *K*_*flag*_ = 2.7 *×* 10^5^*k*_*B*_*Tb*_*x*_, and *T*_*m*_ = 300*k*_*B*_*T* .

#### 3.2.5 Different flagella arrangements

*E. coli* bacteria in nature present a variety of flagella attachment locations ranging from their clustering near one end of the body to a random distribution over the whole body ^36^. Figure 9 shows two *E. coli* models with different arrangements of flagella in comparison to experiments (see also Movies S8-S11). In all simulations, the total number of flagella is five. The symmetric placement of flagella we used so far [see Fig. 9(a)] leads to a persistent run dynamics of the bacterium with a moderate wobbling of the body, as shown in Fig. 3(b). The symmetric model exhibits tumbling that compares well with experimental observations, if the flagella are primarily located near one end of the body (see Movies S8 and S10). We have also tried a model with flagella randomly attached within the rear half of the body, which exhibits a comparable run phase, but does not always reproduce successful tumbling, as the reverted flagellum does not always leave the bundle quickly enough. Finally, a random distribution of flagella attachment points on the whole body results in a strong wobbling of the body [see Fig. 9(b)] with a reduction in the swimming speed. For this model, the tumble phase is also not well controlled, such that the change in the swimming direction ranges from small to large angles (see Movie S9). Similar run-and-tumble dynamics can also be found in experiments, as shown in Fig. 9(b) and Movie S11. Thus, it is difficult to generalize the run-and-tumble behavior of *E. coli* bacteria, when flagella are randomly distributed over the whole body. Interestingly, experimental observations indicate that all types of flagella arrangements are possible, though the majority of bacteria have them clustered near one end of the body. Note that the distribution of flagella attachment points can easily be adopted in simulations to reproduce the variety of *E. coli* tumbling dynamics in experiments.

**Figure 9.**
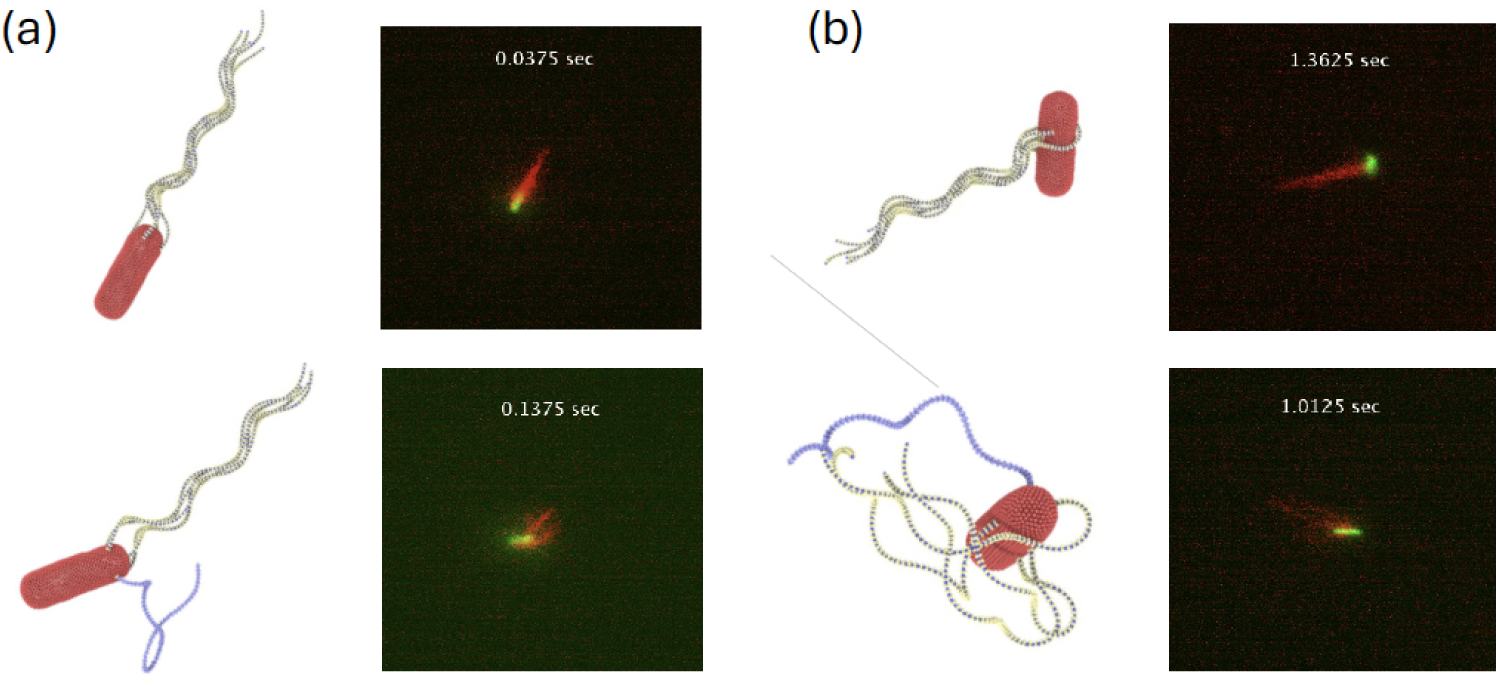
*E. coli* models with different arrangements of flagella in comparison to experiments. (a) A symmetrical arrangement of flagella with one flagellum attached at the back along the body axis and the other four attached symmetrically at the body circumference with a 90^°^ separation. In the experiment, the flagella are primarily located near one end of the body. See also Movies S8 and S10. (b) Random distribution of flagella attachment points on the whole body (see Movies S9 and S11). Snapshots in the top row show the run phase, while the bottow row illustrates the tumble phase. In simulations, *N*_*flag*_ = 5, *K*_*flag*_ = 2.7 *×* 10^5^*k*_*B*_*Tb*_*x*_, *T*_*m*_ = 300*k*_*B*_*T* , and the hook rigidity of the reverted flagellum is 500*k*_*B*_*T* with polymorphic transformation, while the other flagella have *K*_*hook*_ = 100*k*_*B*_*T* .

#### 3.2.6 Tumbling with several reverted flagella

Experimental observations suggest that more than one reverted flagella may be involved in the tumble phase. Figure 10 displays snapshots of tumbling *E. coli* models with one [Fig. 10(a)] and two [Fig. 10(b)] reverted flagella for different *N*_*f lag*_. Even though the tumbling behavior is qualitatively similar in both cases, we can also observe some differences (see Movies S8 and S12-S16). The major difference is that changes in the body and bundle orientations are more pronounced when two reverted flagella are activated, as shown in Fig. 11. This is not entirely surprising, since two reverted flagella exert larger torques on the body, enhancing dynamic changes in its orientation. Figure 10 also presents several experimental snapshots, which closely resemble simulation snapshots.

**Figure 10.**
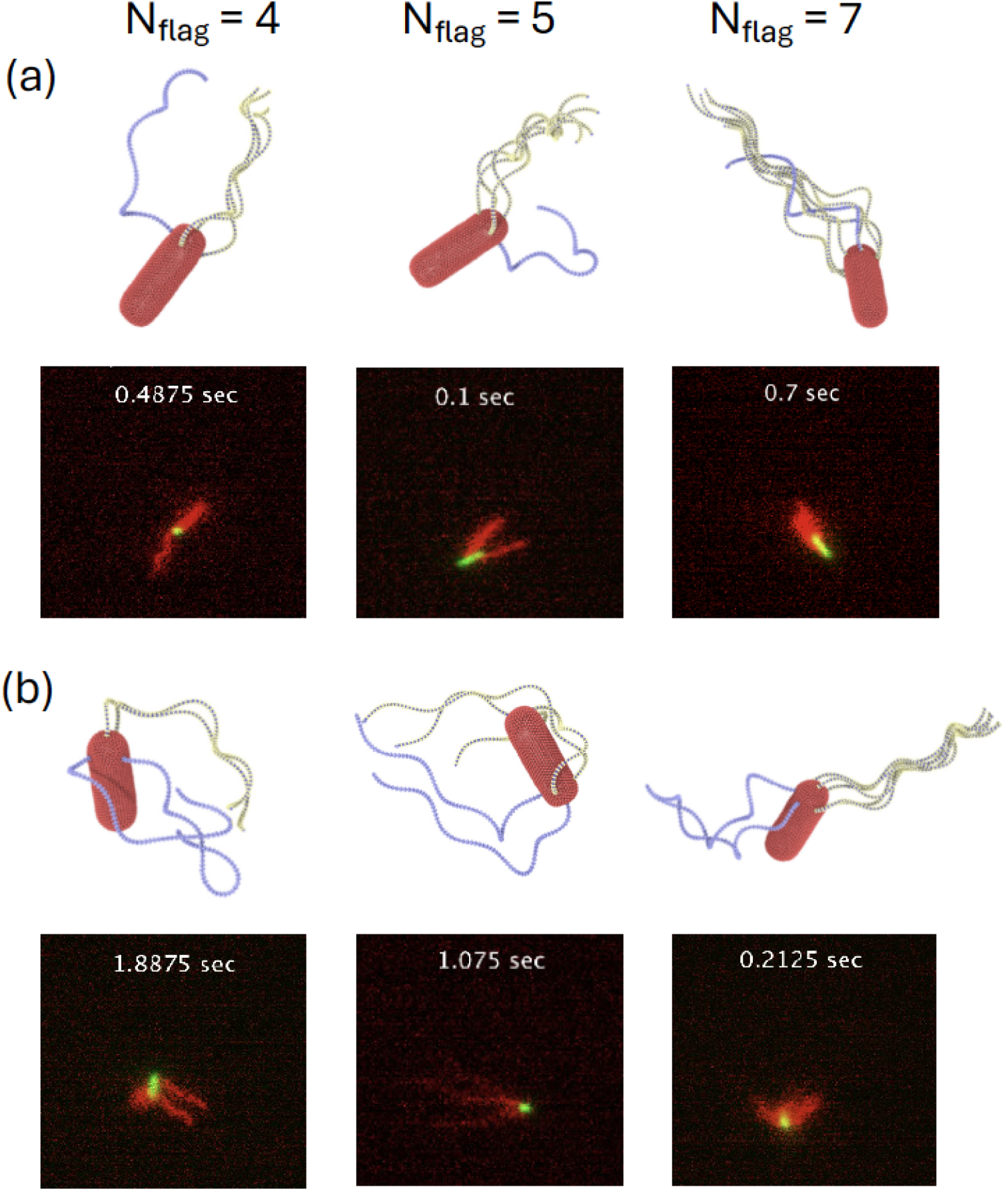
Tumble phase of *E. coli* models with a different number of flagella. Snapshots of several cases with (a) one reverted flagellum, and (b) two reverted flagella. For comparison, several experimental snapshots are also included next to the simulation snapshots. Note that *N*_*flag*_ values should be considered only for the simulation snapshots, as in the experimental observations, it is not possible to count the number of flagella. See also Movies S8 and S12-S16. In all simulations, *K*_*flag*_ = 2.7 *×* 10^5^*k*_*B*_*Tb*_*x*_, *T*_*m*_ = 300*k*_*B*_*T* , and the hook rigidity of the reverted flagellum is 500*k*_*B*_*T* with polymorphic transformation, while the other flagella have *K*_*hook*_ = 100*k*_*B*_*T* .

**Figure 11.**
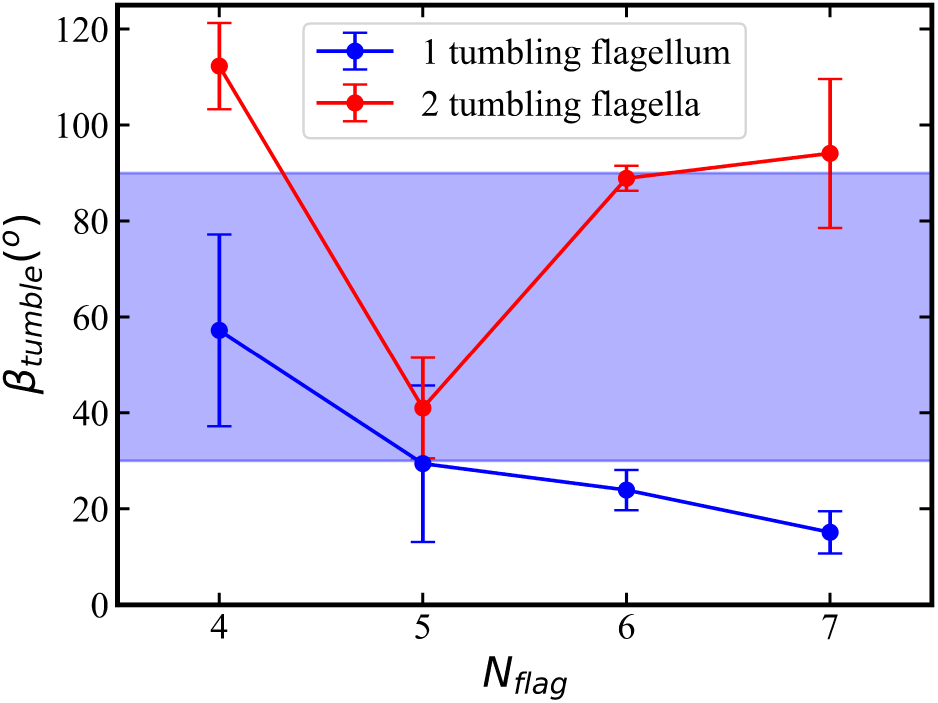
Tumble angles *β*_*tumble*_ of *E. coli* with different numbers of flagella (see snapshots in Fig. 10). The blue curve represents cases with one reverted flagellum, and the red curve corresponds to simulations with two reverted flagella. The blue shade indicates the range of tumble angles measured experimentally ^10^. The tumble angle is defined as the angle between the orientation vector of the body before and after a tumble event.

For a quantitative comparison, we employ the tumble angle *β*_*tumble*_ of *E. coli* shown in Fig. 11 for various *N*_*f lag*_. The tumble angle *β*_*tumble*_ is defined as the angle between the axis of the body before (i.e., in the run phase before tumbling) and after (i.e., when the bundle has formed again) a tumble event. For a single reverted flagellum, *β*_*tumble*_ decreases with increasing *N*_*f lag*_, as it becomes more difficult for the reverted flagellum to leave the bundle [see Fig. 10(a)]. Furthermore, a bundle with more flagella better controls the direction of the body, allowing smaller changes in its orientation. However, when two reverted flagella are involved in tumbling, the behavior of our model during tumbling is more stochastic, leading to larger deviations in *β*_*tumble*_. Furthermore, tumbling angle is affected by the relative position of the reverted flagella on the body. Note that most of the tumble angles in simulations lie within the range of 30 − 90^°^ experimentally observed values ^5,10,24,37^, which is indicated by the blue shaded area in Fig. 11.

Since experimental measurements of flagella bending rigidity yield a rather wide range of 10^−24^ − 10^−21^*Nm*^2 39^, we have also considered the possible effect of *K*^*flag*^ on the tumble angle *β*_*tumble*_, shown in Fig. 12. Simulations indicate that *E. coli* with *K*_*f lag*_ = 1.8 *×* 10^4^*k*^*B*^*Tb*_*x*_ does not exhibit a significant tumble behavior with *β*_*tumble*_ ≈ 0. For larger *K*_*f lag*_, *β*_*tumble*_ first decreases starting from relatively soft flagella with *K*_*f lag*_ = 4.5 *×* 10^4^*k*_*B*_*Tb*_*x*_ to *K*_*f lag*_ = 2.7 *×* 10^5^ *k*_*B*_*Tb*_*x*_. For even larger *K*_*f lag*_, an increase in *β*_*tumble*_ is observed. Note that in the limit of rigid flagella, no bundle formation due to hydrodynamic interactions would be possible ^12^, even for a very flexible hook.

**Figure 12.**
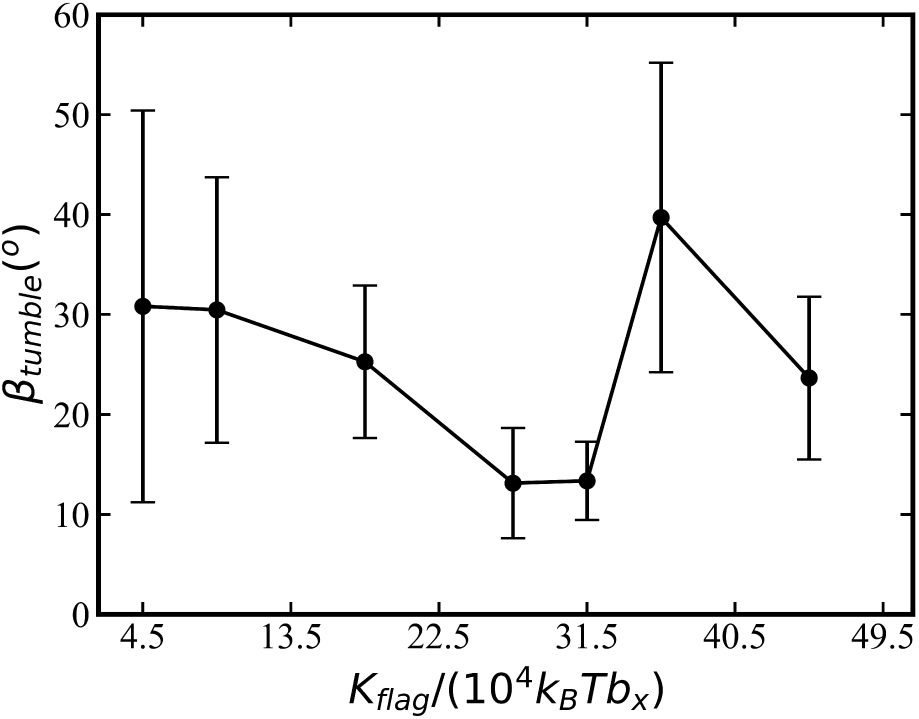
Tumble angle *β*_*tumble*_ of *E. coli* as a function of flagella bending stiffness. Here, only one reverted flagellum is employed with *N*_*flag*_ = 5. The tumble angle is defined as the angle between the orientation vector of the body before and after a tumble event. Each data point is averaged over three tumble events.

## 4 Discussion and conclusions

In this work, our main goals were to develop a realistic and flexible *E. coli* model, and to address the question, which physical properties of *E. coli* govern its run-and-tumble behavior. We have identified a few parameters which strongly affect *E. coli* swimming behavior, including the shape of the body and flagella, polymorphic transformation of flagella, dynamic hook stiffening, and the arrangement of flagella on the body. It is a proper combination of these characteristics that makes *E. coli* an excellent swimmer which can efficiently explore the surrounding environment.

An interesting result from both simulations and experiments is that the swimming speed is nearly independent of the number of flagella. Even though the total actuation power increases with increasing *N*_*f lag*_ in simulations since each flagellum is driven by the same constant torque, the swimming velocity remains nearly unaffected. The rotation frequency of the bundle increases slightly with increasing *N*_*f lag*_, however, the formed bundle appears to be less tight for a large number of flagella. Experimentally measured swimming velocities also appear to be independent of the number of flagella, though it is less clear whether the total actuation power increases with increasing *N*_*f lag*_ or not. A possible explanation for this result can be that the number of flagella mainly affects the thickness of the formed bundle. Since the propulsion force depends logarithmically on the bundle thickness, it only weakly affects the swimming velocity. Furthermore, the existence of *E. coli* with a different number of flagella suggests that there should not be significant evolutionary advantages of a specific number of flagella.

An important aspect is the ratio between rotational frequencies of the body and the flagellar bundle, which is close to 1*/*5 for a wild-type *E. coli*. During the run phase, the rotational frequency of flagellar bundle determines the swimming speed, while rotation of the body plays only a minor role. However, during the tumble phase, the rotational frequency of the body becomes important. When the body rotates fast, flagella that leave the bundle may quickly wrap around the body due to its rotation, limiting significantly the ability of a bacterium to change direction. Therefore, the ratio of 1*/*5 between the body and bundle rotation frequencies is small enough to facilitate the fast rotation of the bundle for a fast swimming speed and not to be detrimental for the tumble phase. For a spheroidal body in Fig. 5, this ratio is close to 1*/*2, implying that the ability for an efficient bacterium tumbling with a not-very-fast rotation of the body imposes a much stronger constraint on the maximum rotation frequency of the bundle, and thus on the maximum swimming speed. Note that at low torques (*T*_*m*_ ≲ 200*k*_*B*_*T* ) of the actuating motors, the formation of a tight flagellar bundle is impeded due to weak hydrodynamic iterations, resulting in a reduced propulsion force of *E. coli*. Therefore, for efficient run-and-tumble behavior, a proper balance between the body and bundle rotation is essential, which depends on geometric characteristics of the body and the bundle, the arrangement of the flagella, and the value of the applied torque.

The other important physical property for an efficient *E. coli* tumbling is the polymorphic transformation of clockwise-rotating flagella that leave the bundle. Tumbling starts when one or several flagella switch the direction of rotation from anti-clockwise to clockwise. If the helicity of the clockwise-rotating flagella does not change, they start propelling against the swimming direction, leading to a substantial slow down of *E. coli* and partial disruption of the tight bundle. As a result, the reverted flagella do not leave the bundle far enough to facilitate significant change in the swimming direction. However, when polymorphic transformation takes place and the reverted flagella change their helicity in addition to the change in the rotation direction, they exert propulsion forces in a direction close to the swimming direction, which does not significantly disturb the bundle. Furthermore, in this case, the reverted flagella and the bundle do not attract each other hydrodynamically, which facilitates the motion of the reverted flagella away from the bundle, resulting in an efficient change of the swimming direction. A further important characteristic for *E. coli* tumbling is the bending rigidity of the hook, which controls the angle between the initial part of flagella and the body surface. For bundle formation during the run phase, flagella hooks have to be not too stiff, so that flagella can bend enough near their base and gather together into the bundle. However, the hook rigidity also plays a vital role in the tumble phase. Forces from a bent hook act to bring flagella to an orientation perpendicular to the body, aiding the reverted flagella to move away from the bundle. In fact, a recent experimental study ^36^ provides evidence that the bending rigidity of the hook is larger when flagella rotate clockwise in comparison to anti-clockwise rotation. This measurement suggests stiffening of the hook of reverted flagella, which has been considered in our simulations and has a positive effect on *E. coli* tumbling. Furthermore, the preference of a perpendicular orientation of flagella with respect to the body surface due to the hook bending rigidity affects its dynamics. Imagine a cable with torsional rigidity mounted per-pendicular to a surface and bent along the surface. Rotation of this cable at the base would result first in pulling it along the surface due to torsional resistance until it bends enough near the base to allow the full rotation at the base. Note that if the mounting angle of the cable is not restricted (i.e., flexible attachment), the cable base would be aligned with the surface and its rotation at the base would simply lead to the rotation of the whole cable without any pulling effect. Similarly, reverted flagella attached to the cylindrical side of the body are first pulled out of the bundle, if the hook is not too soft, facilitating their escape from the bundle. Thus, these two effects (hook stiffening and initial pulling of reverted flagella) provided by the bending rigidity of the hook aid in the successful escape of the reverted flagella from the bundle.

Despite the fact that the developed *E. coli* model properly reproduces experimental observations and helps to explain the importance of different physical properties for bacterium dynamics, it should be best considered as a set of *E. coli* models. Large differences in *E. coli* behavior have been found when different arrangements of flagella are considered. In fact, a substantial variety in run-and-tumble behavior of *E. coli* is also observed in experiments, suggesting that structural properties of *E. coli* may have wide distributions. For our base model, we have selected *N*_*f lag*_ = 5 symmetrically placed flagella at one of the body ends for simplicity. Random placement of flagella near one of the body ends yields a behavior which is quite similar to that with the symmetric placement of flagella. However, random distribution of flagella attachment points at the whole surface of the body results in a run-and-tumble behavior, which strongly differs from that exhibited by the base model. Systematic characterization of *E. coli* behavior for all combinations of parameters is currently not feasible due to high dimensionality of the parameter space. Furthermore, the behavior of *E. coli* should be considered in a statistical fashion, since each tumbling event is different even for the base model with symmetrically placed flagella. For instance, a few tumbling attempts in simulations were not successful, as the reverted flagellum is not able to leave the bundle.

In the future, this model can be applied to some questions related to the behavior of *E. coli* and other peritrichous bacteria in complex environments, where experimental observations may not be detailed enough to propose corresponding physical mechanisms. For example, a recent experimental investigation ^42^ suggests that *E. coli* tumbling is necessary for a bacterium to escape from the wall, while the underlying physical mechanism is not clear. Another interesting case is *E. coli* swimming within lipid vesicles, leading to the formation of thin membrane tethers and overall vesicle transport by the encapsulated bacteria ^58^.

## Acknowledgements

We acknowledge funding from the ETN “Physics of microbial motility” (PHYMOT) within the European Union’s Horizon 2020 research and innovation programme under the Marie Skłodowska-Curie grant agreement No 955910. R.B., A.L. and E.C. acknowledge the ANR-22-CE30 grant-Push-pull. B.W.-Z., G.G. and D.A.F. gratefully appreciate computing time on the supercomputer JU-RECA ^59^ at Forschungszentrum Jülich under grant no. actsys.

